# Accurate detection of complex structural variations using single molecule sequencing

**DOI:** 10.1101/169557

**Authors:** Fritz J. Sedlazeck, Philipp Rescheneder, Moritz Smolka, Han Fang, Maria Nattestad, Arndt von Haeseler, Michael C. Schatz

**Author notes:** Equal contribution (Fritz J. Sedlazeck, Philipp Rescheneder).

## Abstract

Structural variations (SVs) are the largest source of genetic variation, but remain poorly understood because of limited genomics technology. Single molecule long read sequencing from Pacific Biosciences and Oxford Nanopore has the potential to dramatically advance the field, although their high error rates challenge existing methods. Addressing this need, we introduce open-source methods for long read alignment (NGMLR, https://github.com/philres/ngmlr) and SV identification (Sniffles, https://github.com/fritzsedlazeck/Sniffles) that enable unprecedented SV sensitivity and precision, including within repeat-rich regions and of complex nested events that can have significant impact on human disorders. Examining several datasets, including healthy and cancerous human genomes, we discover thousands of novel variants using long reads and categorize systematic errors in short-read approaches. NGMLR and Sniffles are further able to automatically filter false events and operate on low amounts of coverage to address the cost factor that has hindered the application of long reads in clinical and research settings.

## Introduction

Structural variations (SVs), including deletions, duplications, insertions, inversions and translocations of at least 50bp, account for the largest number of divergent base pairs (bp) across human genomes^1^. SVs have been shown to contribute to polymorphic variation, pathogenic conditions, and large-scale chromosome evolution^2^, and several human diseases such as cancer^3^, autism^4^, or Alzheimer’s^5^ have been associated with SVs. SVs have also been shown to impact phenotypes for an increasing number of other organisms^6–10^. Nevertheless, SV calling is still in an early stage with large numbers of false positives (i.e. erroneously calling a SV that is not actually present) and false negatives (i.e. not calling a genuine SV that is present).

One of the first reports of the prevalence and importance of SVs came in 2004, when Sebat, et al.^11^ used microarrays to discover large-scale copy number polymorphisms were surprisingly common across healthy human genomes. Today, SV detection is most commonly performed using short paired-end reads, such as Illumina sequencing, where variances in coverage or paired-end alignments indicate a mutation with respect to a reference genome. Copy number variations can be observed as decreases (deletions) or increases (amplifications) in aligned read coverage^12^, and other types of SVs can be identified by the arrangement of paired-end reads or split-read alignments^13–16^. Short read approaches, however, have been widely reported as lacking sensitivity, with only 10%^17^ to 70%^6,8^ of SVs detected, very high (up to 89%) false positive rates ^6,18–21^ and misinterpreting complex or nested SVs^6,22^.

The advent of long read single molecule sequencing by Pacific Biosciences (PacBio) and Oxford Nanopore has the potential to substantially increase the reliability and resolution of detecting SVs. With read lengths averaging around 10kbp and some reads exceeding 100kbp, the reads can be more confidently aligned to repetitive sequences that often mediate the formation of SVs^22^. In addition, long reads are more likely to span a SV so that the breakpoints can be captured by high-confidence alignments. Furthermore, long reads enable improved phasing, which is necessary to study overall genome structure and allele specific characteristics. Despite these advantages, long single molecule sequencing reads also introduce new challenges. Most significantly, these technologies have a very high sequencing error rate, which is currently around 10% to 15% for PacBio, and 5% to 20% for Oxford Nanopore sequencing^23^. These high error rates exceed the capabilities of most aligners or SV detection algorithms and require new, specialized methods. A few aligners have been proposed, including BlasR^24^, BWA-MEM^25^ and more recently GraphMap^26^. However, only one standalone method, PBHoney^18^, is available to detect all types of SV from long read data, although a few others have been proposed for subset of SVs types such as SMRT-SV^27^ that focuses on insertion, deletions (indels) and inversions.

To address these challenges, we introduce a pair of novel open-source analysis algorithms, NGMLR and Sniffles, for comprehensive long read alignment and SV detection (**Figure 1**). NGMLR (https://github.com/philres/ngmlr) is a fast and accurate aligner for long reads based on our previous seed-and-extend short read aligner NGM^28^, extended with a new segmented convex gap-cost scoring model to align high error long reads across SV breakpoints. Its partner algorithm Sniffles (https://github.com/fritzsedlazeck/Sniffles) successively scans within and between the alignments to identify all types of SVs. Addressing the high error rate of the reads, Sniffles employs a novel SV scoring scheme to exclude false SVs based on the size, position, type and coverage of the candidate SV. This is particularly important for PacBio sequencing where false-positive indels are the predominant error type and Oxford Nanopore sequencing which shows systematic indel errors in homopolymer sequences and other contexts.

**Figure 1:**
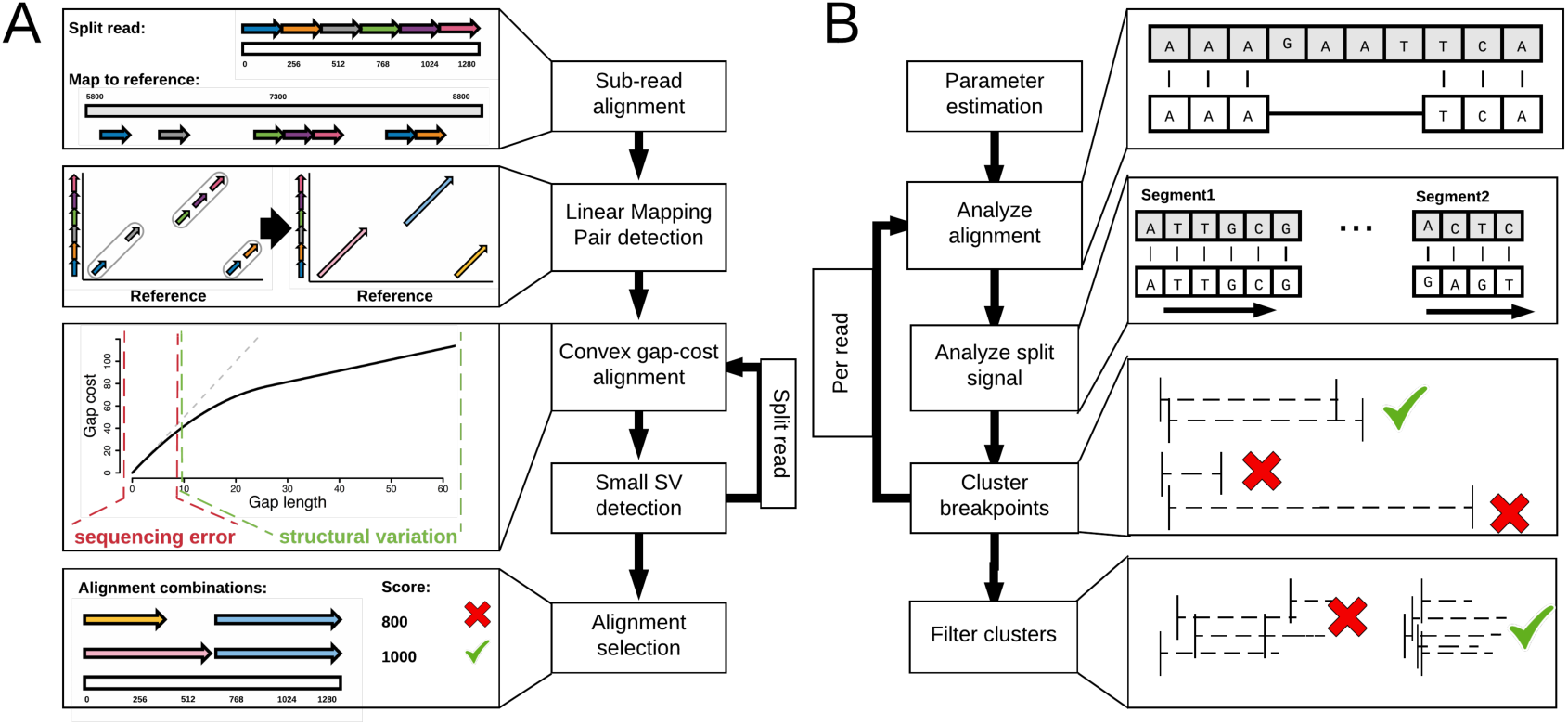
Overview of the main steps implemented in NGMLR (A) and Sniffles (B). For details see Supplementary Sections 1 and 2 for NGMLR and Sniffles, respectively.

We apply our methods to several simulated datasets and genuine datasets of Arabidopsis, healthy human genomes, and a cancerous human genome to demonstrate the increased accuracy compared to alternate short and long read callers. A particularly novel feature of Sniffles is its ability to detect nested SVs, such as inverted tandem duplications (INVDUP) or inversions flanked by insertions or deletions (INVDEL). These are poorly studied classes of SVs, although both have been previously associated to genomic disorders^29–31 32^. However, as no alternative methods can routinely detect them, their full significance is currently unknown. Finally, we show that our methods can reduce the costs per sample in both the sequencing coverage and computational resources required, making it increasingly feasible to apply long read technologies to large numbers of samples.

## Results

### Accurate mapping and detection of SVs using long reads

NGMLR is our novel alignment method to accurately align long, high error reads, even in the presence of SVs (**Figure 1a**). A major innovation of NGMLR is the use of a convex gap scoring model^33^, which allows it to accurately align reads spanning genuine indel SVs in the presence of small indels (1-10bp) that commonly occur as sequencing errors. Larger or more complex SVs are also captured through split-read alignments (**Figure 2**). To achieve both high performance and high accuracy, NGMLR first partitions the long reads into 256bp sub-reads and aligns them independently to the reference genome. It then groups co-linear sub-read alignments into long segments, which are then aligned using a dynamic programming algorithm with our convex gap-cost scoring scheme. Finally, NGMLR selects the highest scoring non-overlapping combination of segments per read and outputs the results in standard SAM/BAM format. See **Methods** and **Supplementary Section 1** for more details.

**Figure 2:**
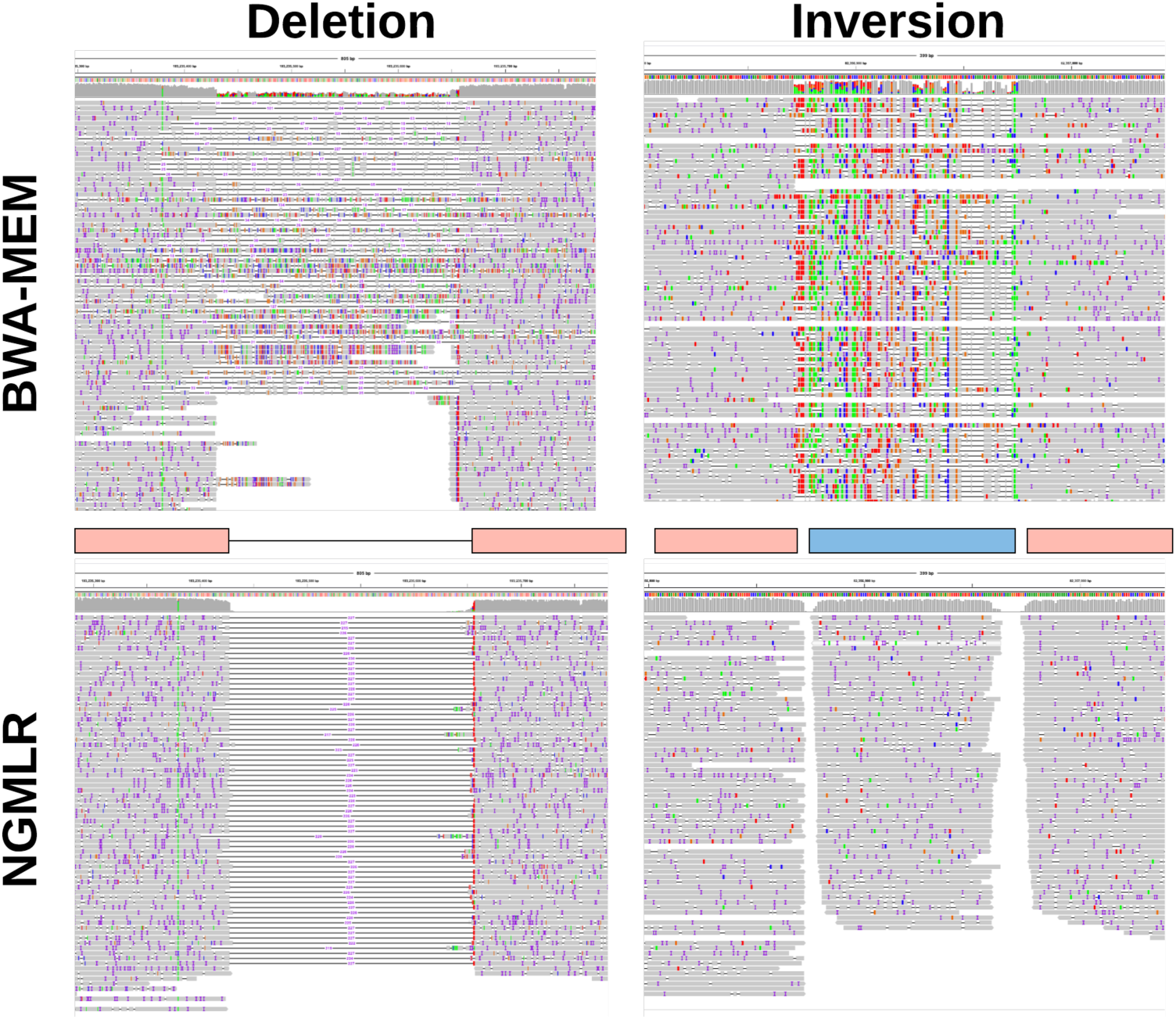
Alignment improvements using NGMLR shown for a 228 bp deletion (left) and a 150 bp inversion (right) shown in IGV^34^. Upper track shows BWA-MEM alignments that indicate these events but is not able to localize the precise event and breakpoints. With the improved alignments of NGMLR, Sniffles can precisely pinpoint the location and type of the SV.

We further present Sniffles to detect all types of SVs (deletions, duplications, insertions, inversions, translocations, and nested events) from long read alignments. It can be used with any aligner, although we find it has the best performance with NGMLR as it produces the most accurate alignments. The principal steps of Sniffles consist of scanning the alignments of each read independently for potential SVs and then to cluster and refine the candidate SVs across all reads (**Figure 1b**). Sniffles uses both within-alignment and split-read information to detect SVs, as small indels can be spanned by a single alignment, but large or complex events lead to split-read alignments. A major innovation of Sniffles is the novel SV scoring it uses to filter false SV signals from the noisy PacBio and Oxford Nanopore reads. Like other variant detectors, filtering by minimum read support (default: 10 reads) is a critical feature, but it also considers new features such as the consistency of the breakpoint position or size. In addition, Sniffles can perform read-based phasing of SVs and report adjacent or nested events in the output VCF file. See **Methods** and **Supplementary Section 2** for more details.

#### Evaluation of NGMLR on simulated human data

Using the error profiles and read lengths measured from two human datasets (**Supplementary Section 3.2**), we simulated 50x PacBio-like and 50x Oxford Nanopore-like read data sets from two human chromosomes (chr21 and chr22). In the simulation, we included a total of 840 SVs consisting of equal numbers of indels, duplications, balanced translocations, and inversions ranging from 100bp to 50kbp in size (**Methods**). **Figure 3a** summarizes the results when evaluating NGMLR, BWA-MEM, BLASR and GraphMap aligning these reads to the entire human genome^24–26^. Each bar represents one data set consisting of 20 SVs of a certain type and length, and categorizing the read alignments as: *precisely* capturing the breakpoints and the correct type of the SV (green); *indicating* the type but without exact break points (yellow); *trimmed* so that the region of the read containing the SVs was not aligned (gray); *forced*, such as the BWA-MEM alignments in **Figure 2** (red); *fragmented* so that a read is split more often than necessary (brown); or the entire read was *unaligned* (white) (**Methods** and **Supplementary Table 1**). Across all SV types, NGMLR outperforms the other mappers with an average 80.32% precisely aligned versus 26.31% for BWA-MEM as the next closest competitor. Even when counting the precise and the indicated representation together, NGMLR outperforms with an average 91.83% versus 69.17% for BWA-MEM as next closest competitor.

**Figure 3:**
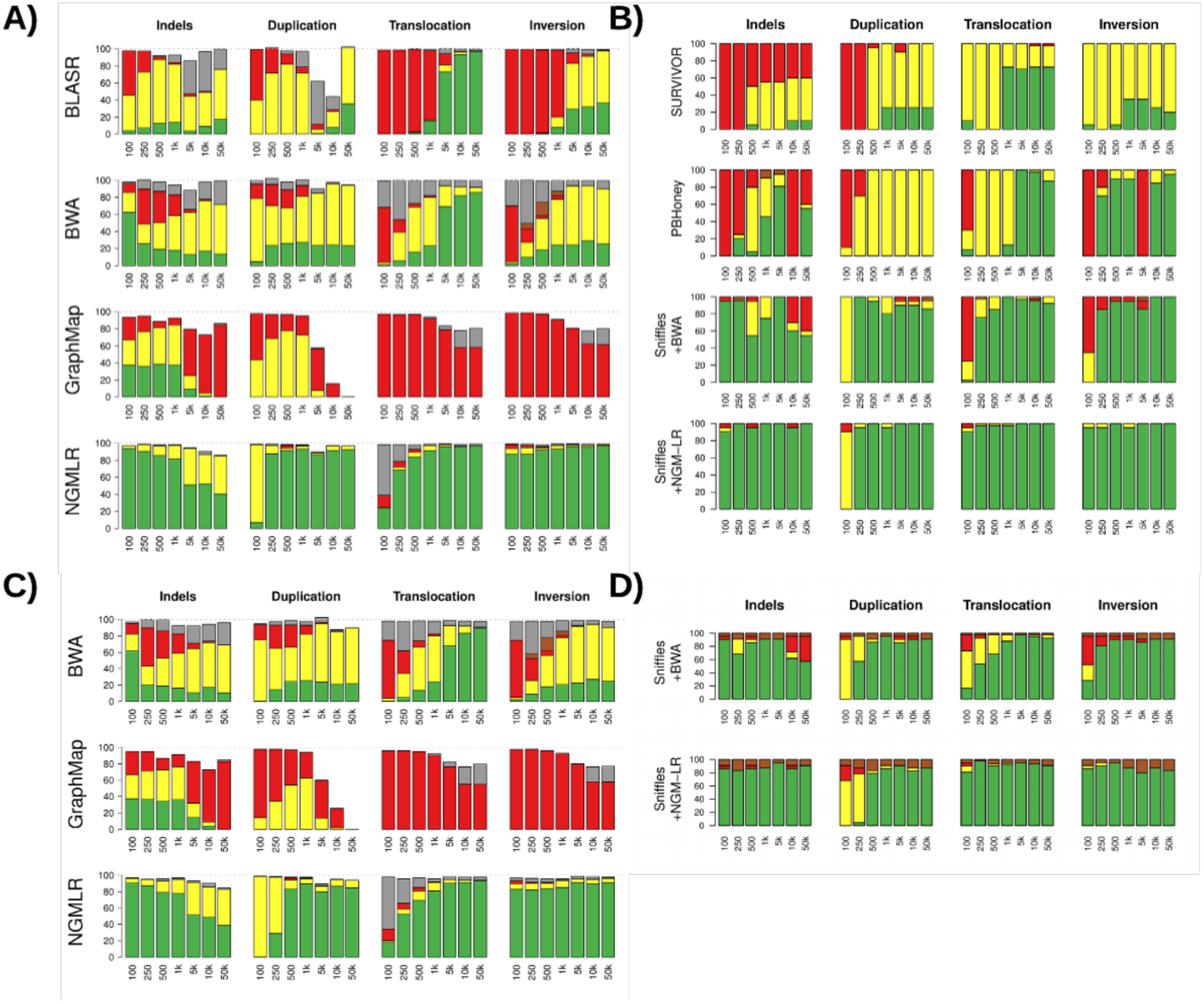
Evaluation of NGMLR and Sniffles using simulated data with 840 SVs. For read alignments, we simulated PacBio-like (a) and Oxford Nanopore-like reads (c), and distinguish between: *precise* (green), *indicated* (yellow), *forced* (red), *unaligned* reads (white), or *trimmed* but not aligned through the SV (grey). The SV analysis (b,d) used the same alignments as before, and distinguishes between: *precise* (green), *indicated* (yellow), *not indicated* (red) and *false positive* calls (brown).

Next, we compared the performance of NGMLR, BWA-MEM and GraphMap in mapping simulated Oxford Nanopore-like reads, using their respective parameter suggestions (BlasR was excluded, as it is only applicable to PacBio reads). Again, NGMLR substantially outperformed other mappers for *precisely* aligning reads (72.42% vs 24.90% for the second best BWA-MEM), or when considering both *precise* and *indicating* alignments (88.57% versus 67.96% for BWA-MEM) (**Figure 3c**). GraphMap performed rather poorly on these data, with on average only 18.19% of reads aligned *precisely* or *indicating* the SV as it *forces* 61.13% the reads to align across the SV.

#### Evaluation of Sniffles based on simulated human data

Next, we evaluate the performance of Sniffles compared to alternate short and long read SV detection approaches using the alignments reported above ^14–16,18^ (**Figure 3b**). We also extended the analysis to include simulated short reads to be analyzed by our consensus algorithm SURVIVOR^8^. SURVIVOR aggregates the outputs from Lumpy, Manta and Delly and excludes variants reported by only a single caller. We find this increases specificity without sacrificing much sensitivity8. Similar to the read alignments, we classified SVs to be: *precisely* detected if they are reported within +/-10bp (green); *indicated* if they are within +/-1kbp and ignoring the type (yellow); *not detected* (red); and *false positive* (brown) (**Methods** and **Supplementary Table 2**).

Over all SV types, the combination of Sniffles and NGMLR performs the best with an average of 94.20% *precisely* detected SVs and an FDR of 0.00%. The most problematic class was short (100bp) tandem duplications, as they are identified as insertions rather than tandem duplications, and hence classified as *indicated*. The second best result was achieved using Sniffles with BWA-MEM alignments, with on average 79.11% *precisely* detected SVs and a 0.68% FDR. With the more noisy BWA-MEM alignments, Sniffles detects the presence of an SV, but cannot reliably predict the exact location or sometimes even the type of SV. For example, both deletions and inversions cause an accumulation of mismatches in the BWA-MEM alignments (**Figure 2**). PBHoney, which relies on BlasR alignments, *precisely* detected only 32.58% of simulated SVs and missed 25.18%. Most of the 40.73% *indicated* SVs from PBHoney came from misinterpreting tandem duplications as insertions. For the short-read analysis, SURVIOR *precisely* detected 18.81% and 57.89% *indicated* of the simulated SVs, similar to what has been previously reported for short read analysis^6,8^, although the consensus-based analysis reduced the FDR to 0.17%.

Finally, we benchmarked the performance of Sniffles using BWA-MEM and NGMLR on the Oxford Nanopore-like reads described above (**Figure 3d**). Using Sniffles with NGMLR, 82.25% of SVs are *precisely* identified, whereas 76.35% are *precisely* identified with BWA-MEM. Nevertheless, due to the higher rate of sequencing errors in the Oxford Nanopore-like data, Sniffles using either aligner has a slight FDR of calling 1-4 additional events per data set.

#### Benchmarking NGMLR and Sniffles with genuine long human reads

The simulated read results establish a baseline of performance, although may not capture the full complexity of real sequencing data. To benchmark more realistic datasets, we next analyzed genuine PacBio^35^ and Oxford Nanopore^36^ reads from the well-studied NA12878 human genome. Since a complete truth set of SVs is not available for this genome, we modified the reference human genome to introduce 700 SVs at random locations: 140 insertions (by deleting from the reference), 140 deletions (by adding new sequence), 140 inversions, and 140 balanced translocations creating 280 translocation events. The mean indel and inversion size was 1.6kb. We did not attempt to simulate tandem duplications, as this would require detecting and modifying tandem duplications preexisting in the reference.

In this analysis, we can only evaluate the sensitivity of alignments, but not false positives since there are additional true SVs in the sample compared to the reference. NGMLR showed a clear improvement over BWA-MEM (58.65% vs 32.35%) for *precisely* aligned reads across the SVs (**Supplementary Table 3**), although the shorter average length of the genuine reads limited the number of reads that could be *precisely* aligned. For example, if an insertion is longer than the read length, then the read can only *indicate* the SV. When counting *precise* and *indicated* together, NGMLR achieved a substantially better result than BWA-MEM (76.96% vs 49.21%). Furthermore, NGMLR considerably reduced the number of *forced aligned* reads compared to BWA-MEM (3.01% vs 24.21%). Using the Oxford Nanopore reads from NA12878 we observe a similar trend with NGMLR giving the most *precise* alignments (51.56% vs. 27.35%) with the lowest percent of *forced* reads (5.94% vs. 29.15%).

Using these alignments and the alignment of 50x coverage of genuine Illumina sequencing from this sample^6^, we next benchmarked the available SV callers (**Supplementary Table 4**). Sniffles and NGMLR achieved the highest rate of *precisely* called SVs with 95.14% and 88.29% SVs using the PacBio and Oxford Nanopore reads, respectively. In contrast, the short read-based SURVIVOR analysis had a much lower sensitivity (76.57%) considering either *precise* or *indicated* variants.

### Trio-based analysis of Structural Variations

#### Assessment based on PacBio sequencing of an Arabidopsis trio

We next analyzed an Arabidopsis trio (Col-0, CVI and the Col-0 x CVI F1 progeny) previously sequenced using both PacBio and Illumina sequencing^37^. This is a particularly challenging dataset as the rate of heterozygosity in the F1 is approximately 1 SNP every 200bp along with thousands of reported SVs ^37^. Using Sniffles with default parameters, we identified 355 (Col-0) and 9,652 (CVI) SVs in the parents (**Table 1**), of which 42 (Col-0) and 6,679 (CVI) were homozygous. Based on Mendelian inheritance, we expected all homozygous SVs identified in the parental cultivars to be in the F1 as heterozygous variants. Indeed, when comparing the homozygous calls from Col-0 to the F1 only 4 SVs were not identified. On closer inspection, one missed insertion was reported as 47bp in F1 vs. 53bp in Col-0, and similarly a deletion was reported as 48bp in F1 vs. 53bp in Col-0. Both of these events were initially not found due to the minimum size cutoff of 50bp. Sniffles can detect the remaining two SVs – another deletion and a duplication – in the F1 by reducing the coverage threshold as the deletion was supported by only 4 reads and the duplication by only 3 reads.

**Table 1:** Summary of detected SVs across 15 different data sets. SVs were reported with a min. size of 50bp using SURVIVOR based on Delly, Lumpy and Manta for Illumina or Sniffles for PacBio or Oxford Nanopore requiring at least 10 reads. Supplementary Table 5 shows all the data sets used.

| Data Set | Tech. | Cov. | Avg. read length(bp) | Total SVs | DEL | DUP | INS | INV | TRA |
| --- | --- | --- | --- | --- | --- | --- | --- | --- | --- |
| Arabidopsis Col-0 | PacBio | 127x | 6,482 | 355 | 67 | 63 | 106 | 68 | 51 |
| Arabidopsis CVI | PacBio | 123x | 6,073 | 9,652 | 3,822 | 904 | 1,823 | 478 | 2,625 |
| Arabidopsis Col-0 x CVI F1 | PacBio | 155x | 11,206 | 11,935 | 4,974 | 582 | 4,049 | 567 | 1,763 |
| Arabidopsis Col-0 X CVI F1 | Illumina | 40x | 250 | 10,950 | 4,324 | 643 | 0 | 671 | 5,312 |
| Giab HG002 (son) | PacBio | 69x | 8,540 | 19,131 | 7,957 | 1,084 | 9,656 | 232 | 202 |
| Giab HG002 (son) | Illumina | 80x | 148 | 10,822 | 5,018 | 863 | 0 | 823 | 4,118 |
| Giab HG003 (father) | PacBio | 32x | 6,284 | 11,964 | 5,296 | 408 | 6,048 | 99 | 113 |
| Giab HG003 (father) | Illumina | 80x | 148 | 11,395 | 5,553 | 869 | 0 | 818 | 4,155 |
| Giab HG004 (mother) | PacBio | 30x | 7,285 | 10,463 | 4,590 | 276 | 5,436 | 93 | 68 |
| Giab HG004 (mother) | Illumina | 80x | 148 | 8,901 | 5,000 | 868 | 0 | 829 | 2,204 |
| NA12878 (healthy female) | PacBio | 55x | 4,334 | 15,499 | 6,734 | 606 | 7,880 | 160 | 119 |
| NA12878 (healthy female) | Oxford Nanopore | 28x | 6,432 | 17,155 | 12,301 | 323 | 4,401 | 87 | 43 |
| NA12878 (healthy female) | Illumina | 50x | 101 | 7,275 | 3,744 | 553 | 0 | 731 | 2,247 |
| SKBR3 (Breast Cancer) | PacBio | 69x | 9,872 | 19,165 | 7,268 | 1,019 | 10,391 | 328 | 159 |
| SKBR3 (Breast Cancer) | Illumina | 25x | 101 | 5,046 | 2,776 | 483 | 0 | 627 | 1,160 |

When comparing CVI to the F1 calls, Sniffles initially missed 370 (5.54%) SVs that were reported in CVI and not in the F1. Most of the missed variants are explained by a few straightforward explanations: 159 lacked sufficient coverage of supporting reads in the F1; 101 had slightly different SV sizes reported below the minimum size; 43 were interpreted as different SV types; and 50 occurred within Col-0 specific regions in F1 (**Supplementary Section 4.4**). After considering these factors, only 17 (0.25%) SVs present in the CVI data set were missed by Sniffles for the F1 data set. In contrast, the Illumina-based SURVIVOR calls in the F1 data set had a much lower recall rate compared to the PacBio-based Sniffles in Col-0 (47.3% recall) and CVI (70.6% recall).

#### Genome-in-a-Bottle (GiaB) Human Trio Analysis

Next, we investigated the performance of Sniffles based on the Ashkenazi trio data set from GiaB^38^ (**Table 1 and Supplementary Table 6**). Similar to Arabidopsis, we analyzed the concordance of Mendelian inheritance between samples as an indicator of performance, although some SVs (e.g. mobile element insertions in the son) may be incorrectly classified. We adjusted the coverage threshold for Sniffles to a minimum of 5 reads (-s 5) to account for the reduced coverage of the parents compared to the son (32x compared to 69x, also see downsampling results below). We compared these results to the Illumina-based call sets from 80x coverage in all of the samples.

Sniffles reported 5,244 and 5,964 SVs as homozygous in the father and mother, respectively. Within the son we re-identified 93.84% and 94.01% of the SVs from the father and the mother, respectively. Most of the missed variants could be explained through minor adjustments in parameters. For example, when we relax the size cutoff to consider variants just below 50bp, Sniffles misses only 187 (3.57%) and 126 (2.11%) for the father and mother, respectively, and most of the remainders have slightly less coverage than our cutoff. In contrast, when using SURVIVOR, we identified only 1,586 and 1,668 homozygous SVs for father and mother, respectively, approximately 3 times less than found using Sniffles. Of these, 164 (10.34%) and 203 (12.17%) could not be identified in the son.

We next tested how many calls are in the son that are not within the parents to investigate potential false positive calls (**Supplementary Table 6**). Using the same parameter settings, Sniffles had the lowest number of such calls in the son for deletions (515 vs. 677), inversions (66 vs. 75) and translocations (90 vs. 1,550) compared to SURVIVOR. Only for tandem duplications SURVIVOR has 75 events that are unique to the son versus 115 that Sniffles calls. On investigation, most of the Sniffles calls found only in the son were due to the lower coverage of the parents

Overall, Sniffles and NGMLR had the highest recall rate as well as the lowest Mendelian discordance rate. In contrast, the short read approaches showed an unreasonably high number (1,550) of false positive translocations in the son.

### Comparison of PacBio and Oxford Nanopore sequencing for human SV analysis

As a new sequencing technology, Oxford Nanopore has not yet been extensively tested for structural variation analysis, especially in human genomes. Here we investigated its capability in the well-studied NA12878 human genome using three publicly available datasets: 28x coverage of Oxford Nanopore data^36^ analyzed with NGMLR/Sniffles, 55x coverage of PacBio data^35^ analyzed with NGMLR/Sniffles, and 50x coverage Illumina data^39^ analyzed by SURVIVOR (**Table 1**). We also compared these results to two previously published call sets, the GiaB insertion and deletion call set based on PacBio sequencing^35^ and the Illumina-based deletion-only call set from the 1000 Genomes Project (1KGP)^6^.

We first used this data set to measure the runtime of the different aligners and Sniffles. For this we subsampled the reads to 1x coverage and mapped them with each of the mappers. NGMLR was the fastest requiring 1.3 hours using 10 threads followed by BWA-MEM (1.7 hours). GraphMap did not finish within a week on the same data set and computer using 10 threads. Sniffles required 3.4 hours and 2.2 hours for calling SVs based on the 55x NGMLR PacBio and 28x NGMLR Oxford Nanopore mapping from NGMLR (**Supplementary Table 10**).

Overall, Sniffles identified 15,499 SVs for the PacBio reads, and 17,155 SVs for the Oxford Nanopore reads, while SURIVOR reported 7,275 (Table 1). Sniffles using either PacBio (1,298) or Oxford Nanopore (1,269) reads showed the largest overlap with the 1KGP deletions (**Supplementary Table 7**). Sniffles also recalls the most deletions from the published GiaB NA12878 call set using PacBio (6,641) and Oxford Nanopore (6,557) data, compared to only 3,009 for the Illumina data. For insertions, Sniffles recalls 7,488 and 6,234 of the GiaB results when using PacBio or Oxford Nanopore, respectively, while the short read consensus calls had none.

The majority (25,100 distinct calls) of the identified SVs are present in only one call set, while 16,277 SVs were identified in two or more call sets. However, most (94.38%) of the PacBio calls were confirmed by Oxford Nanopore, Illumina or the existing call sets. Surprisingly, Oxford Nanopore had substantially worse concordance, as Sniffles reports 11,989 calls unique to Oxford Nanopore, of which 11,394 (96.19%) were deletions and the majority (89.72%) were within a homopolymer or other simple repeats. In contrast, the 847 calls only found by PacBio were mainly insertions (68.12%) and only 352 (41.46%) were overlapping with homopolymers or repeats given the same criteria. This systematic bias for deletions in the Oxford Nanopore data is most likely an error in the base calling, as also reported by Jain *et al.* ^29^. The majority of these artifacts are small deletions, and by increasing the minimum SV size to 200bp, Sniffles reports only 38.86% of the SVs calls within homopolymers and low complexity regions. The Illumina-based SV calling had relatively low concordance to alternative approaches, and 31.43% of their calls were unique to the technology. Interestingly, the majority (53.89%) of the unique calls were again translocations events, but most of these appear to be false positives (see below).

### Detailed investigation of unique short read vs. long read events

Over all data sets, Sniffles is able to detect far more indels than the short read based callers (**Table 1**). Conversely, using the short read approach we detect, on average, 27 times more translocation events compared to using Sniffles within presumably healthy human data sets. We investigated these discrepancies using NA12878.

We first investigated the small insertion (50bp-300bp) and deletion (50bp-3kbp) calls from Sniffles using the orthogonal Illumina reads as evidence (**Supplementary Section 4.5.1**). We focused on these size ranges since they should be best captured by the paired-end Illumina data. To do so, we used a derivative of the compression-expansion statistic^40^ as an unbiased measure of the Illumina paired-end placements near predicted indels. Briefly, we compare the genome-wide observed Illumina insert size (average 311bp) to the insert sizes spanning the indel breakpoints as aligned using BWA-MEM: real insertions in the sample cause the pairs to map closer than expected, deletions further away. Using the Illumina data and a p-value threshold of 0.01 (two sided, one sample t-test), we could confirm 3,415 and 3,879 deletions reported by Sniffles in the PacBio and Oxford Nanopore data, respectively (**Supplementary Table 9**). For insertions, we could validate 2,685 and 1,703 for PacBio and Oxford Nanopore, respectively. For comparison, using SURVIVOR we could confirm 1,873 deletions. Manta performed best over the individual short read based callers with 2,868 confirmed deletions and 629 confirmed insertions, although far less than using Sniffles with either long read technology. We assessed the performance of this test by shuffling the regions with the same sizes across the genome and performing the test again. On average only 10% of the randomized regions showed a significant alteration.

Next, we investigated the large number of translocations reported in the Illumina-based consensus calls (2,247) compared to Sniffles (PacBio: 119 and Oxford Nanopore: 43). Note that the number of Illumina-based translocations is even higher before computing the consensus: 3,117 for Delly, 3,007 for Lumpy, and 3,261 for Manta. (**Supplementary Section 4.5.2 and Supplementary Table 9**) We noted a large overlap (48.87%) of the Illumina-based translocation sites with insertion calls from Sniffles using both long read technologies. **Figure 4a** shows a representative example, with an insertion called using both long read data types exactly at the location where the candidate translocation is called. As the insertion falls within a low-complexity region, it causes the short reads to be mis-mapped to a similar repeat on a different chromosome. These mis-mapped reads are therefore incorrectly identified as a translocation instead of an insertion. We also observed clusters of soft-clipped Illumina reads (displayed as red endings of the reads), which further indicates an insertion rather than a translocation event. Overall, we could rule out 1,869 (83.18%) of the Illumina-based translocation calls as false, with most overlapping an insertion (48.87%) or deletion (8.86%), or a few other SV types (1.20%). The remaining Illumina-based translocation calls are also questionable, with 404 (17.98%) being in low complexity regions and 141 (6.28%) translocations lying within a region with abnormally high coverage without any evidence for them in the long read data. A similar pattern is also seen for the inversions, where 60% of the calls overlap with a different SV type identified by long reads (**Figure 4b**) or align to low complexity sequences.

**Figure 4:**
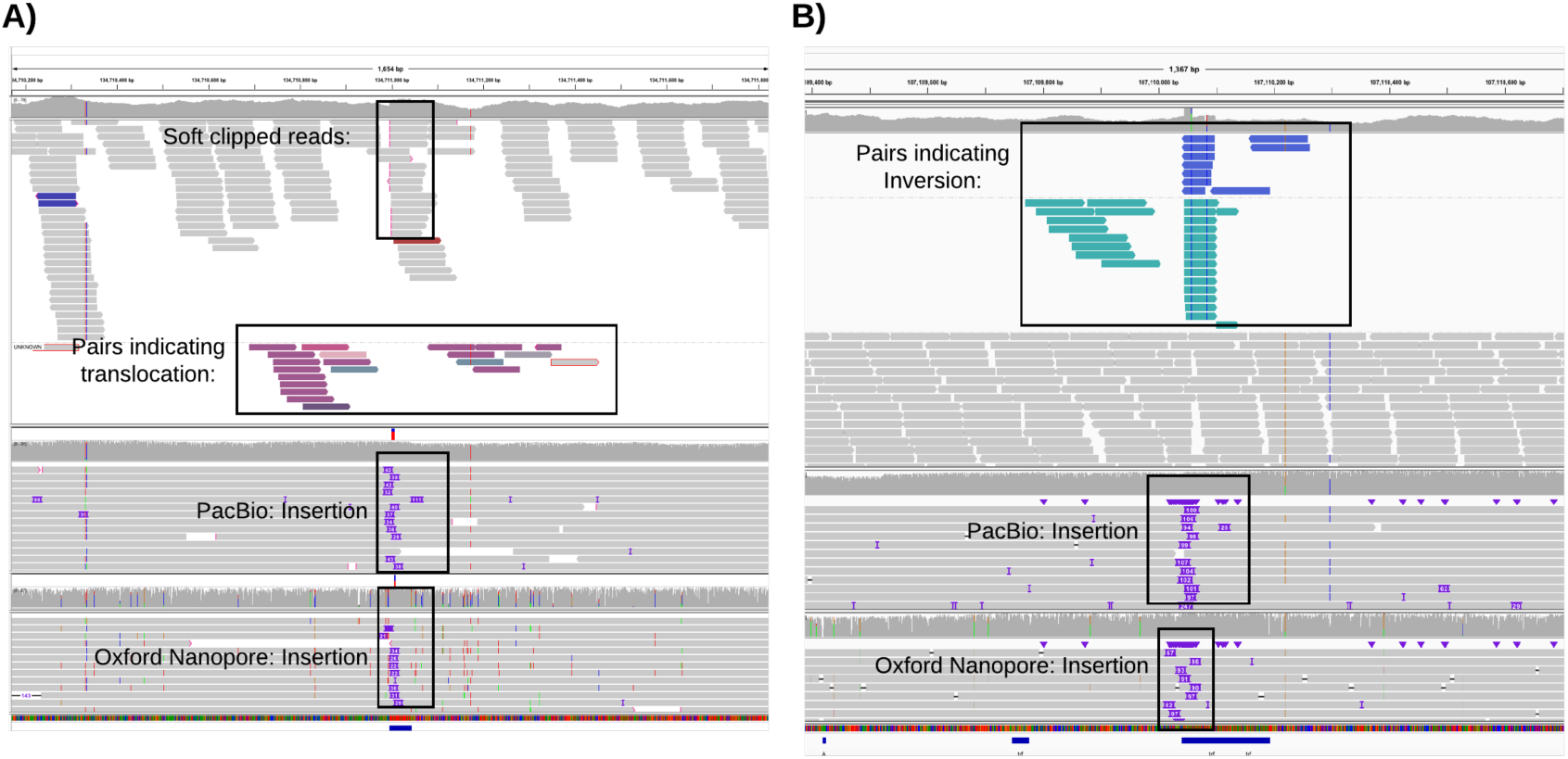
Systematic error in short read based SV calling. A) An example of a putative translocation identified in the short read data (top alignments) that overlaps an insertion detected by both PacBio (middle) and Oxford Nanopore sequencing (bottom). B) An example of a putative inversion identified in the short read data (top) that overlaps an insertion detected by both PacBio (middle) and Oxford Nanopore reads (bottom)

Overall, the majority of PacBio-based insertions and deletions calls from Sniffles were validated by either the Oxford Nanopore calls or the Illumina paired-end reads. In contrast, the majority of calls unique to the Illumina-based methods were false, with false translocations caused by mis-mapped reads intersecting insertions being especially prominent.

### Detection of Nested SVs

Next, we investigated the performance of Sniffles on more complex, nested SV types such as inverted duplications (INVDUP) and inversions flanked by deletions (INVDEL). While these variant types are poorly studied, they have been associated with a number of diseases: INVDUPs have been reported in PLP1 (associated with Pelizaeus-Merzbacher disease ^30^) and MECP2 region^29^ and shown impact on the VIPR2 gene^41^. INVDELs have been reported in Haemophilia A genetic deficiency using long range PCRs^32^.

To start, we simulated nested SV types of different sizes (280 SVs total) in the human genome along with simulated PacBio-like, Oxford Nanopore-like, and Illumina-like reads (**Figure 5** and **Supplementary Table 2**). We evaluated each SV separately e.g. an inversion flanked by two deletions was evaluated based on all three SV. For the short read based caller we observe a reduced ability to detect these complex events, and none of the approaches were able to identify the correct breakpoints of the inversion flanked by deletions. Nevertheless, nearly all methods predicted an inversion to be present. For the inverted duplications SURVIVOR correctly predicted almost all the inversions, but missed the fact that the region is duplicated. We observed a similar result for PBHoney using PacBio-like reads where it called the inversion in 0.00% and 5.32% *precisely* for INVDEL and INVDUP, respectively. Only Sniffles was able to detect the full types due to its dynamic splitting of events, and *precisely* called 67.88% of the nested SVs (**Supplementary Section 2**). This includes SVs that are larger than the read length and thus highlights the ability of Sniffles to accurately infer such complex events. When using the Oxford Nanopore-like reads, our ability for *precisely* calling these events is slightly reduced due to the more complex sequencing error model and reduced read length (average of 6kbp). However, we were still able to *precisely* call 67.34% of SVs on average over INVDEL and INVDUP events of different length.

**Figure 5:**
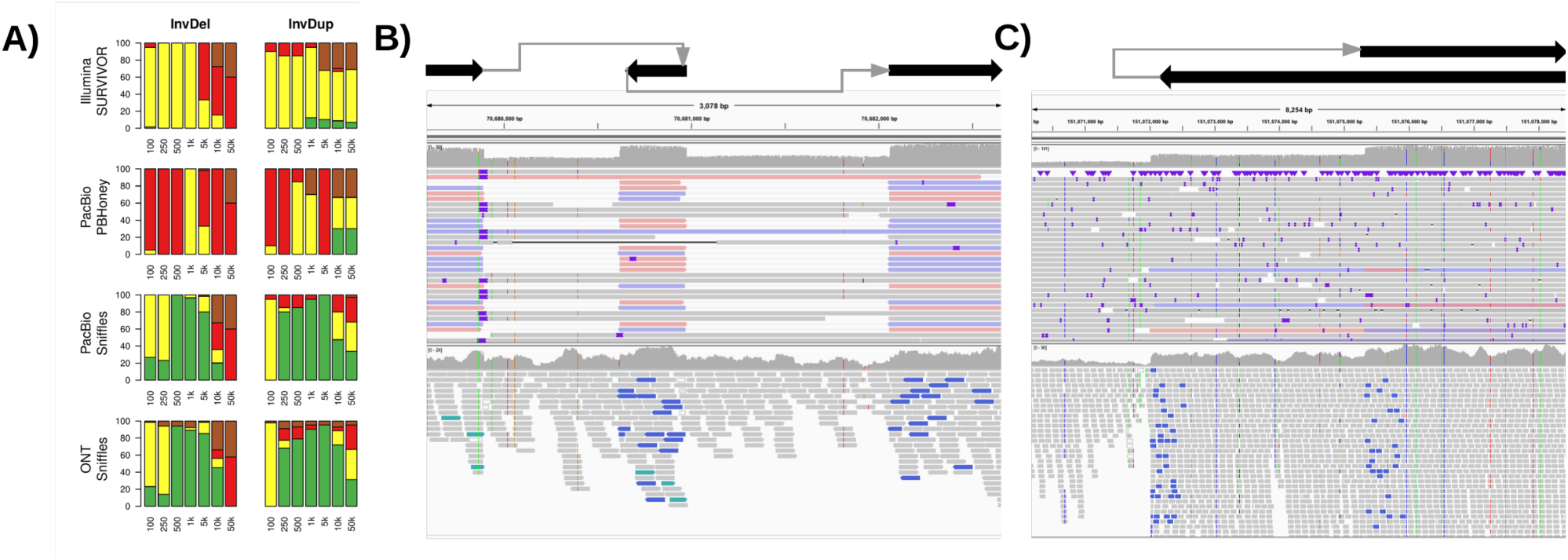
Nested SVs in SKBR3 cancer cell line. A: Evaluation of Sniffles + NGMLR using simulated data to identify nested SVs. B: A 3kb region including two deletions flanking an inverted sequence clearly visible and detected by Sniffles using NGMLR (above) and not detected by the Illumina methods (below). C: The start of an inverted duplication. The breakpoints were reported by Sniffles as the start of an inverted duplication (above) and not correctly detected by short read methods (below).

To highlight this capability in real data, we examined a PacBio-based data set for the SKBR breast cancer cell line (Nattestad et al, in submission). Sniffles and NGMLR was used to investigate this data set revealing 15 gene fusions joined by 1 to 3 chained events, which were all validated by PCR. **Figure 5** shows an INVDEL (B) and INVDUP (C) in SKBR3 in comparison to Illumina short read data (lower panel). The short reads indicate an inversion (colored reads) but the poor resolution makes it impossible to detect and interpret the entire event. In contrast, Sniffles detects the events, and the read phasing allows for the complex regions to be fully resolved (**Supplementary Section 2**). Although these were the only two nested types we evaluated, Sniffles is capable to detect and report multiple combination of SVs based on the IDs assigned in the reported VCF file.

### How much coverage is required?

Our final analysis is to assess how much coverage is required to detect SVs using PacBio or Oxford Nanopore reads. This is an important consideration since long read technologies are more expensive than short read technologies in generating the same amount of coverage^23^. From a purely statistical analysis, assuming Poisson coverage and that an SV can be detected by reads that span 50bp, about 10x coverage should be sufficient to infer all SV breakpoints using 10kbp long reads whereas about 25x coverage is needed for 2x100bp short reads (**Figure 6a** and **Methods**). However, this analysis represents an idealized case (e.g. lack of repeats or coverage biases) and is a weak estimate for the true amount of coverage required.

**Figure 6:**
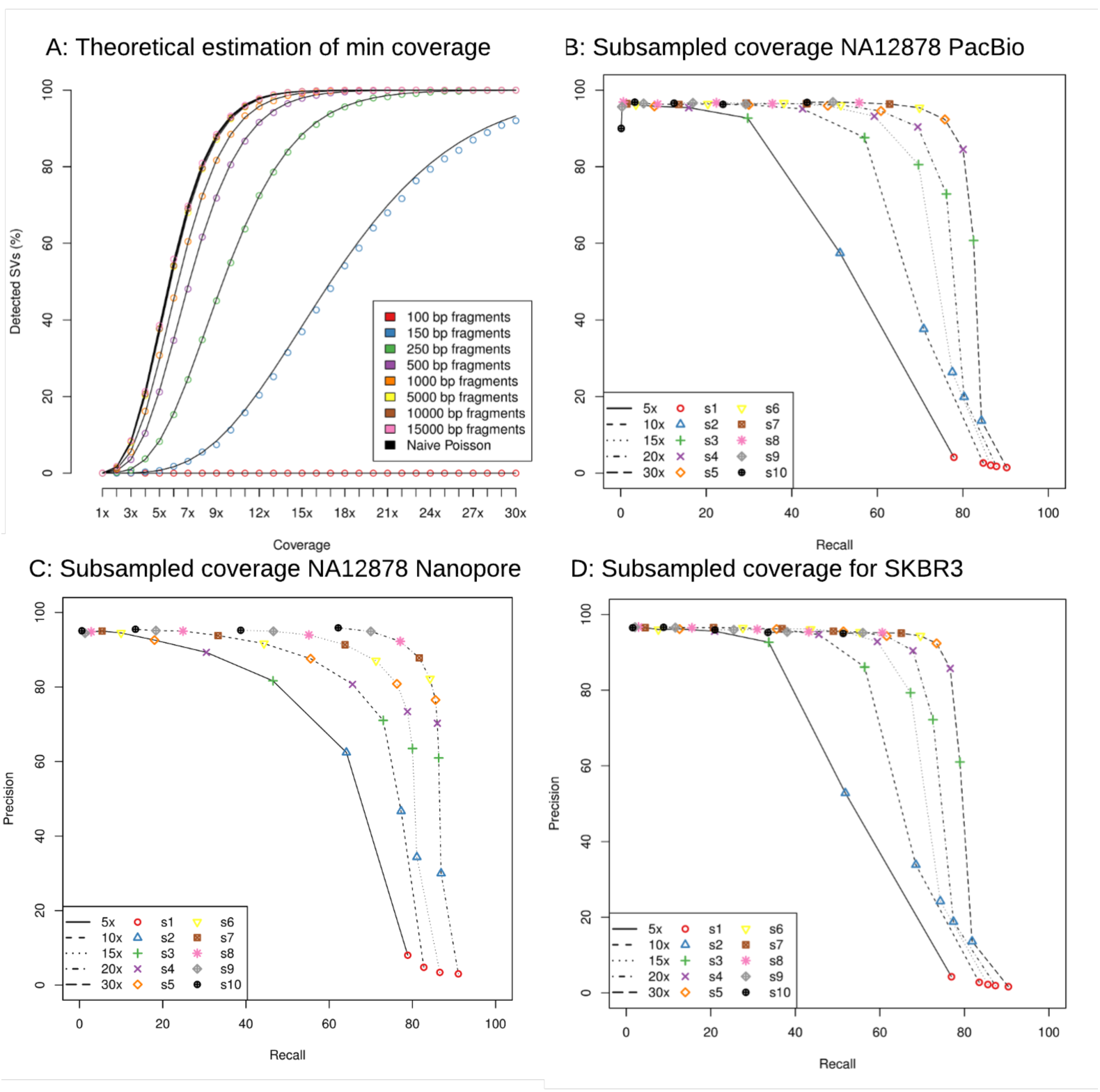
Analysis of SV detection accuracy with different amounts of coverage. A: Theoretical assessment of recall vs coverage for different read lengths requiring a 50bp overlap of each breakpoints for SV events. B: Subsampling experiment of the 55x PacBio NA12878 data; C: Subsampling experiment using 28x Oxford Nanopore NA12878 data; D: Subsampling experiment of the 70x PacBio SKBR3 breast cancer cell line dataset. For plots B-D, Sniffles and NGMLR were run on subsampled data (rate indicated by lines) and using different thresholds for Sniffles (s: 1-10 indicated in symbols and colors). In every data set we could show the success for Sniffles using NGMLR with only 10x to 30x coverage that recovers around 80% of the calls with a precision ~80% or higher.

To investigate this with real data, we subsampled reads from the NA12878 PacBio and Oxford Nanopore datasets as well as the more complex SKBR3 PacBio sample to 5x, 10x, 15x, 20x and 30x coverage. We then aligned those reads using NGMLR and used Sniffles with different parameter settings (-s 1 to –s 10 to vary the minimum number of reads) to compare the results to the original call set. For this analysis, we measured precision and recall with respect to the full coverage dataset (**Figure 6b-d**). As expected, using a minimum support of only one or two reads (red dots, blue triangles) to support an SV leads to many false positives. Consequently, we focus on those settings that have a precision rate of 80% or higher.

For both PacBio datasets (**Figure 6 b,d**) we obtain similar results that show 15x coverage has a precision of ~80% and recall of 69.64% and 67.24% for NA12878 and SKBR3 for homozygous and heterozygous SVs of any type, respectively. The difference in recall is largely due to the complexity of the SKBR3 cancer sample, which displays high copy (>20 fold) amplifications in different regions of the genome. Increasing the coverage to 30x, Sniffles has an 80.05% to 76.63% recall with a precision of ~85% for NA12878 and SKBR3, respectively.

For the Oxford Nanopore NA12878 data set, the highest recall rate (84.23%) had a precision of 82.24% for 20x coverage (**Figure 6 c**). The higher apparent accuracy in this case is not too surprising, since the original data set has only 28x coverage, so this constitutes a less dramatic down sampling. Interestingly, we see a greater loss in precision than the PacBio data, due to the stringent minimum number of supporting reads (-s 10) used throughout the study.

Overall, we could demonstrate that using NGMLR and Sniffles we can detect the vast majority of heterozygous and homozygous SVs using only a fraction of the original coverage.

## Discussion

In this paper, we introduced NGMLR, a novel long read mapper, and Sniffles, a novel long read based SV caller. The versatility of these methods enables an unprecedented view into structural variations in the human genome and other genomes from long read single molecule sequencing data. We demonstrated their reliability over several simulated and genuine data sets, where the new methods outperformed existing tools in sensitivity and specificity. In particular, by leveraging long reads we demonstrated that we can overcome the sensitivity issues reported for short read callers, which can miss between 30%^6,8^ and 90%^17^ of the SVs. This allows us to detect tens of thousands of additional variants beyond what has been reported by large-scale short-read sequencing projects such as the 1000 Genomes Project. Furthermore, prototype versions of our methods were used in a recent study to identify the causal, pathogenic SV in a patient who presented with multiple neoplasia and cardiac myxomata^42^. We could also use the long read data to identify systematic errors in short read structural variation analysis, where the vast majority (>85%) of the translocations are false positives due to mis-mapped reads.

The assessment of SVs from long reads is a challenging problem. Errors and artifacts can originate anywhere in the process, including in sequencing, base calling, alignment, or in SV calling. While the predominate error mode for PacBio sequencing is insertions or deletions of a few bases, we have discovered that it also introduces larger false insertions at a low, but noticeable rate (**Supplementary Section 2.2.4**). We have controlled for this artifact by requiring the size and composition of candidate SVs are consistent across the spanning reads, although this artifact limits accuracy in very low coverage data sets. Within the Oxford Nanopore dataset, we have highlighted major artifacts in base calling that form systematic artificial deletions in low complexity repeats. Consequently, they appear as genuine variants to the SV caller, and while we fully expect this to improve in time, it is currently necessary to exclude small SVs when using Nanopore sequencing. Beyond sequencing errors, we highlighted how various alignment artifacts can lead to miscalling or missing of SVs. For example, some long read mappers align reads through a SV without indication of the underlying events. Although Sniffles is able to recognize such errors through the increase in mismatches in the alignment, it is clear that using NGMLR is a better choice to accurately align the long reads. Finally, we showed a deficiency in detecting nested variations such as INVDUP or INVDEL in all examined methods except for Sniffles. While several diseases are already known to be associated with these SV types, we expect their importance will rapidly increase as more samples are analyzed using our methods.

The last remaining barrier to perform a detailed analysis of SVs across a large number of samples is cost. Long read technologies from both PacBio and Oxford Nanopore are becoming less expensive every year, but still remain more expensive than short read sequencing^23^. We have addressed this by investigating how much coverage is needed for accurate SV calling, and have shown how high accuracy is possible with only 15x to 30x coverage for a healthy or cancerous human genome. This translates to a potential price reduction of several tens of thousands of dollars per sample. These requirements will be reduced even more in the years to come as the throughput and read length increase and sequencing error rates decrease. Altogether, these improvements, aided by our methods, will usher in a new era of high quality genome sequences for a broad range of research and clinical applications, and lead to new insights into polymorphic variation, pathogenic conditions, and the forces of evolution.

## Online Methods

### NGMLR

NGMLR is designed to accurately map long single molecule sequencing reads from either Pacific Biosciences or Oxford Nanopore to a reference genome with the goal of enabling precise structural variation calls. We follow the terminology used by the SAM specification ^43^ where a read mapping consists either of one linear alignment covering the full read length or multiple linear alignments covering non-overlapping segments of the read (i.e. split reads).

The main challenge when mapping high error long reads is to evaluate whether a read should be mapped to the reference genome with one linear alignment, or must be split. For example, the correct mapping for a read that spans an inversion can only be found when splitting the read into three segments. Reads that do not span a structural variation should always be mapped with a single linear alignment. However, error rates are high, and are not always uniform with some regions having an error rate of over 30%. These segments can cause read mappers to falsely split a read. Furthermore, the high insertion and deletion sequencing error of long read technologies cause current read aligners to falsely split up large SVs into several smaller ones and make it difficult to detect exact break points.

To address these challenges, NGMLR implements the following workflow (**Figure 1a**):

1. NGMLR identifies sub-segments of the read and of the reference genome that show high similarity and can be aligned with a single linear alignment. These segments can contain small insertions and deletions, but must not span a larger structural variation breakpoint. In reference to BLAST’s High-scoring Segment Pairs (HSPs), we call those segment linear mapping pairs (LMPs).
2. For each LMP, NGMLR extracts the read sequence and the reference sequence and uses the Smith-Waterman algorithm to compute a pairwise sequence alignment using a convex gap cost model that accounts for sequencing error and SVs at the same time.
3. NGMLR scans the sequence alignments for regions of low sequence identity to identify small SVs that were missed in step (1) and (3).
4. Finally, NGMLR selects the set of linear alignments with the highest joint score, computes a mapping quality for each alignment and reports them as the final read mapping in a SAM/BAM file.

#### Convex scoring model

When aligning high error long-reads it is crucial to choose an appropriate gap model as there are two sources of insertions and deletions (indels). Sequencing error predominantly causes very short randomly distributed indels (1-5bp) while longer indels (20bp+) are caused by genomic structural variations. A linear or affine gap model appropriately models indels originating from sequencing error, but cannot account for longer indels from genomic variation as these large blocks occur as a single unit, not as the combination of multiple single base insertions or deletions. However, for noisy long reads, the gap-open penalty falsely causes short indels from sequencing error to cluster together and has only little effect on longer indels, especially in repetitive regions of the genome. With the convex scoring model of NGMLR, extending an indel is penalized proportionally less the longer the indel is. Therefore, the convex scoring model encourages large alignment gaps, such as those occurring from a structural variation, to be grouped together into contiguous stretches (extending a large indel has relatively low cost), while small indels, which commonly occur as sequencing errors, to remain separate (extending a 1 bp gap has almost the same cost as opening a new gap).

Using a convex gap model to compute optimal alignments increases computation time substantially as each cell in the alignment matrix not only depends on three other cells, but on the full row and column it is located in ^33^. This would make it infeasible to use convex gap costs for aligning large long-read datasets, so we adapted a heuristic implementation of the convex gap cost algorithm found in the swalign library (https://github.com/mbreese/swalign). Instead of scanning the full cell and row while filling the alignment matrix, we use two additional matrixes to store indel length estimations for each cell. Furthermore, we use the initial sub-segment alignments to identify the part of the alignment matrix that is most likely to contain the correct alignment and skip all other cells of the matrix during alignment computation. (**Supplement Section 1**).

### Sniffles

Sniffles operates within and between the long read alignments to infer SVs. It applies five major steps (**Figure 1b**).

1. Sniffles first estimates the parameters to adapt itself to the underlying data set, such as the distribution in alignment scores and distances between indels and mismatches on the read, as well as the ratios of the best and second best alignments scores.
2. Sniffles then scans the read alignments and segments to determine if they potentially represent SVs.
3. Putative SVs are clustered and scored based on the number of supporting reads, the type and length of the SV, consistency of the SV composition, and other features.
4. Sniffles optionally genotypes the variant calls to identify homozygous or heterozygous SVs.
5. Sniffles optionally provides a clustering of SVs based on the overlap with the same reads, especially to detect nested variants.

For details on each step see **Supplementary Section 2**. In the following, we focus on the methods that are unique to Sniffles, which are the detection and analysis of alignment artifacts to reduce falsely called variants and the clustering of variants.

#### Putative Variant Scoring

The high error rate of the long reads induces many alignments that falsely appear as SVs. Sniffles addresses these by scoring each putative variant using several characteristics that we have determined to be the most relevant to detecting SVs. The two main user thresholds are the number of high quality reads supporting the variant (set using the –s parameter) as well as the standard deviation of the coordinates in the start and stop breakpoint across all supporting reads. The minimum variant size reported defaults to 50bp, but can also be adjusted using the –l parameter. To account for additional noise in the data and imprecision of the breakpoints we use a quantile filtering to ignore outliers given a coverage of more than 8 reads. The computed standard deviations for both breakpoints are compared to the standard deviation of a uniform distribution representing spurious SV breakpoints reported in the region. SVs are only reported if both breakpoints are below this threshold. If the standard deviation for both breakpoints is < 5bp, the coordinates are marked as PRECISE in the VCF file. See **Supplement Section 2.2.4.**

#### Variant Scoring and Genotyping

At the start of the program the user may specify that the VCF output should be genotyped. In this case, Sniffles stores summary information (coordinates and orientation) about all high quality reads that do not include a SVs in a binary file. This includes those reads that support the reference sequence that pass the thresholds for MQ and alignment score ratio. After the detection of SVs, the VCF file is read in, and Sniffles constructs a self-balancing tree of the variants. With this information, Sniffles then computes the fraction of reads that support each variant versus those that support the reference. Variants below the minimum allele frequency (default: below 30%) are considered unreliable; variants with high allele frequency (default: above 80%) are considered homozygous; and variants with an intermediate allele frequency are considered heterozygous. Note Sniffles does not currently consider higher ploidy, however this will be the focus of future work. See **Supplement Section 2.3.**

#### Clustering and Nested SVs

To enable the study of closely positioned or nested SVs, Sniffles optionally clusters SVs that are supported by the same set of reads. Note that Sniffles does not fully phase the haplotypes, as it does not consider SNPs or small indels, but rather identifies SVs that occur together. If this option is enabled, Sniffles stores the names of each read that supports a SVs in a hash table keyed by the read name, with the list of SVs associated with that read name as the value. The hash table is used to find reads that span more than one event, and later to cluster reads that span the one or more of the same variants. In this way Sniffles can cluster two or more events, even if the distance between the events is larger than the read length. Future work will include a full phasing of hapolotypes including SVs, SNPs and other small variants. See **Supplement Section 2.4.**

### Mapping and SV Evaluation

#### Simulation of SV and reads

As described above, SVs were simulated on chromosome 21 and 22 of the human genome (GRCh37). Data sets were generated with 20 variants for each type of SV (tandem duplication, indel, inversion, translocation and nested) and sizes of these events (100, 250, 500, 1kb, 5kb, 10kb and 50kb). Illumina reads were simulated as 100bp paired end reads using the default parameters of dwgsim. For Pacbio and Oxford Nanopore we scanned the alignments of HG002 (GiaB) and NA12878, respectively, and measured their error profile using SURIVOR (option 2). The error profiles and read lengths were used to simulate the reads for each simulated SV data set (**Supplementary Section 3.2**).

#### Modified reference analysis

To allow for a more realistic scenario, we also modified the human reference (GRCh37) and used real reads to assess the introduced SVs. Here we could only simulate a subset of SV types to be insertions, deletions, inversions and translocations. We simulated 140 variants of each type on the human genome (GRCh37) using SURVIVOR (option 1).

#### Software Versions and Parameter settings

BWA-MEM (version 0.7.12-r1039) ^25^ was used with “-M” parameter to map the short reads and with “-X pacbio -M” to follow the recommended parameter settings for PacBio reads. The parameter –M is used to mark only one alignment as primary and the subsequent alignments as secondary. BlasR (version 1.3.1)^24^ was run using the parameters “-sam - bestn 1 -nproc 15” to obtain only the best alignment in SAM format using 15 threads. Furthermore, Blasr was run with the parameters suggested by PBHoney ^18^ “-nproc 15 - bestn 1 -sam -clipping subread -affineAlign -noSplitSubreads -nCandidates 20 - minPctIdentity 75 -sdpTupleSize 6”. SAMTools (version 0.1.19-44428cd) ^43^ was used to convert the SAM alignment files to BAM and to sort the aligned reads.

Delly (version v0.7.3) ^15^, Lumpy (version 0.2.13) ^14^ and Manta (version 1.0.3) ^16^ were used to call SVs over the Illumina reads followed by SURVIVOR (version 0.0.1) ^44^ to combine the calls and report the consensus variants. To allow for the uncertainty with short read variant positioning, SVs were considered to be the same if their start and stop coordinates fell within 1kb of another and were of the same type. PBHoney (version PBSuite_15.8.24)^18^ with default parameters was used to infer SV based on the specified BlasR alignments. The output was converted into a VCF using SURVIVOR (option 10).

#### Evaluation of long read mappings

All simulated reads were mapped to the human reference genome (GRCh37) using BWA-mem ^25^, BLASR ^24^, GraphMap ^26^ and NGMLR. Reads that overlap or map in close proximity to a simulated SV were extracted from the BAM files and used for evaluation. For the genuine datasets, we first mapped the reads to the unmodified reference genome (without SV) using BWA-MEM and extracted all reads that would span our simulated SV by at least 500 bp. Only these reads were then mapped to the modified reference genome using the four read mappers and used for evaluation.

Both simulated and genuine reads were then divided into six categories (**Supplementary Figure 3.5**):

1. Read mappings are considered “*precise*” if they fully identify the SV they cover. To fall into this category, read mappings have to cover all parts of the SV that are required for identification, e.g. a read mapping to an inversion has to cover the inverted part of the genome as well as the non-inverted sequences flanking the inversion. Furthermore, correct mappings have to be split at the simulated breakpoints (+/-10bp) of the SV.
2. Read mappings are considered *“indicated”* if they show the presence of the correct SV but as the wrong type, e.g. a duplication that is represented as an insertion, or show the correct SV but do not show the exact borders (>10bp away).
3. Read mappings are considered “*forced*” if they indicated the wrong number of SVs (e.g. several small instead of a single long insertion) or contain a significant portion of mapping artifacts (eg. not simulated mismatches) over > 10% of the SV length. These include mappings such as a read that is aligned through an deletion or inversion (**Figure 2, top**).
4. Read mappings are considered “*trimmed*” if they have been soft-clipped or otherwise trimmed so that they cannot indicate the SV and do not contain randomly aligned base pairs (ie. noisy regions)
5. Read mappings that are split into more parts than required to cover the underlying SV are classified as “*fragmented*”.
6. Read mappings that are supposed to map across the SV but are not mapped are considered “*unaligned*”.

For all simulated SV types and sizes and all mappers, we count how many reads fall into the above categories, normalize by the number of simulated reads and visualize the result as barplots.

#### Evaluation of SV callers

After the SV calling each VCF file was evaluated using SURVIVOR ^44^ with appropriate parameter sets to compare the variants to the truth set. A SV is considered *precise* if its start and stop coordinate is within 10bp of the simulated start and stop coordinate and the caller predicted the correct type. A SV is considered *indicated* if the start and stop coordinate of the SV is within +-1kb of the simulated event regardless of the inferred type of SV. A simulated SV is considered *not detected* if there is no call that fulfill the two previous criteria. A SV is considered *false*-*positive* if the event was not simulated.

## Acknowledgements

We would like to thank W. Richard McCombie, Sarah Wheelan, Sara Goodwin, and Bui Quang Minh for helpful discussions. This work was supported through National Science Foundation awards (DBI-1350041, IOS-1732253, and IOS-1445025) and National Institutes of Health award (R01-HG006677). P.R. acknowledges support by the RNA-DK Biology (W1207-B09)

